# Exploring virus-host interactions through combined proteomic approaches identifies BANF1 as a new essential factor for African Swine Fever Virus

**DOI:** 10.1101/2024.12.05.627126

**Authors:** Juliette Dupré, Katarzyna Magdalena Dolata, Gang Pei, Aidin Molouki, Lynnette C Goatley, Richard Küchler, Timothy K Soh, Jens B Bosse, Aurore Fablet, Mireille Le Dimna, Grégory Karadjian, Edouard Hirchaud, Christopher L Netherton, Linda K Dixon, Ana Luisa Reis, Damien Vitour, Marie-Frédérique Le Potier, Axel Karger, Grégory Caignard

## Abstract

African swine fever virus (ASFV) causes a highly lethal disease in pigs and represents a significant threat to the global pork industry due to the lack of effective vaccines or treatments. Despite intensive research, many ASFV proteins remain uncharacterized. This study aimed to elucidate the functions of two ASFV proteins, MGF360-21R and A151R, through comprehensive analysis of their interactions with host proteins. Using affinity purification-mass spectrometry and yeast two-hybrid screening approaches, we identified the host protein barrier-to-autointegration factor 1 (BANF1) as a key interactor of both viral proteins. Biochemical and colocalization assays confirmed these interactions and demonstrated that MGF360-21R and A151R expression leads to cytoplasmic relocalization of BANF1. Functionally, BANF1 silencing significantly reduced ASFV replication, indicating its proviral role. Given BANF1’s established function in regulating the cGAS/STING-dependent type I interferon (IFN-I) response, we postulated that A151R and MGF360-21R could inhibit this pathway. Using different strategies, we showed that both A151R and MGF360-21R did indeed inhibit IFN-I induction. Generation of ASFV deficient of A151R or MGF360-21R showed that both mutant viruses enhanced the host IFN response in primary porcine macrophages compared to wild-type virus. However, their capacity to inhibit this pathway could occur through mechanisms independent of BANF1. Proteomic analysis of BANF1 interactors during ASFV infection highlighted potentially roles in chromatin remodeling, nuclear transport, and innate immune response pathways. Altogether, our data provide new insights into ASFV-host interactions, identifying BANF1 as an important new host factor required for replication and uncovering novel functions for A151R and MGF360-21R.

**Author Summary:** African swine fever virus (ASFV) is a highly contagious and deadly disease affecting pigs worldwide, for which there are currently no effective vaccines or treatments. Despite extensive research, many ASFV proteins remain poorly understood. Our study investigated two ASFV proteins, MGF360-21R and A151R, to better understand their functions and interactions with host proteins. Using proteomic approaches, we found both these viral proteins interact with a host protein called barrier-to-autointegration factor 1 (BANF1). Importantly, BANF1 silencing significantly reduced ASFV replication, indicating its important role in the viral life cycle. We also showed that MGF360-21R and A151R help the virus evade the immune system by blocking the production of interferons, which are key defensive molecules against viral infections. However, this immune evasion does not seem to depend on their interaction with BANF1. Additionally, our analysis of BANF1’s interactions during ASFV infection revealed potential roles in chromatin remodeling, nuclear transport, and the innate immune response. These findings provide new insights into how ASFV interacts with its host and highlight BANF1 as a critical factor in viral replication and immune evasion. Our work contributes to a better understanding of ASFV and could pave the way for developing more effective strategies to fight this virus.

## Introduction

African swine fever virus (ASFV) is a large enveloped virus classified within the *Asfarviridae* family. It is the causative agent of African swine fever (ASF), a disease that exclusively affects suids. Transmission occurs through direct animal-to-animal contact, ingestion of contaminated meat products, fomites, and via soft ticks of the genus *Ornithodoros*. ASFV represents a significant risk to pig health and food security globally due to the absence of available vaccines or treatments, and is a disease notifiable to the World Organisation for Animal Health (WOAH). The high lethality in domestic pigs also makes ASFV a significant threat to the global pig industry and imposes a substantial socio-economic burden on many countries. Its double-strand DNA genome, ranging from 170 to 193 kb in size, contains a relatively conserved, evolutionarily stable “core region” at its center flanked by regions that are dominated by multigene family (MGF) genes. Infection of mammalian cells with ASFV triggers the expression of over a hundred viral gene products (1–3) that can extensively interact with the cellular pathways of the host (4). Although ASFV exhibits genomic resemblance with other large DNA viruses, such as poxviruses and herpesviruses, nearly half of ASFV genes lack any known or predictable function. Intriguingly, MGFs and their predicted protein products do not share significant similarities with other known genes or proteins.

Five distinct MGFs (MGF-100, −110, −300, −360, and −505/530) have evolved through gene duplication by homologous recombination (5). ASFV strains lacking or exhibiting only low virulence in pigs are characterized by gene deletions within MGF360 and MGF505 (6,7). The naturally attenuated OUR T1988/3 strain, isolated from a tick during the epizootic situation in Portugal at the end of the 20th century (8), is distinguished from virulent strains by the absence of eight genes: MGF360-10L, −11L, −12L, −13L, −14L and MGF505-1R, −2R, and −3R (9,10). Other studies have shown that MGF genes can determine the host range (11), ASFV virulence (6,12), and survival of infected macrophages (13). Notably, MGF360 proteins can affect the host antiviral immune response by inhibiting type I IFN (IFN-α/β) production (14–17), and deleting individual genes can result in attenuation of ASFV virulence (15) as well as a reduction of virus replication in ticks (11). However, the functions of most of these genes remain unknown. We are still missing a comprehensive view of the interactome that MGF360 proteins establish with the host proteome, although such information is instrumental to better understanding the ASFV replication cycle and pathogenesis at the molecular level.

To further advance in this direction, we utilized a combination of two proteomics methodologies, affinity tag purification-mass spectrometry (AP-MS) and high-throughput yeast two-hybrid (HT-Y2H), to systematically search for cellular interactors of ASFV proteins. This study was initiated by examining interacting partners of the so far uncharacterized ASFV protein MGF360-21R. First, we used an AP-MS approach in cells infected with ASFV to identify both host and viral partners of MGF360-21R. This led us to characterize a dense virus-host interactions network around MGF360-21R, including 232 cellular proteins and one single ASFV protein, A151R. While the role of A151R in virus replication (18) and virulence *in vivo* (19) is evident, the precise molecular mechanisms underlying its functions remain uncertain. Therefore, using the same AP-MS approach with A151R as a bait, we identified 33 specific interactors of A151R, but more interestingly, 48 partners shared with MGF360-21R, among which BANF1 (Barrier-to-autointegration factor 1, also called BAF) emerged as the main interactor.

BANF1 contributes to multiple cellular processes, including post-mitotic assembly (20–24), nuclear membrane repair (25,26), DNA damage response pathway (27,28), and the recruitment of transcription factors (27,29,30). Its functions seem to be associated with maintaining genome integrity (31). More recently, BANF1 has also been shown to play a role in the modulation of the IFN-α/β pathway by acting on the cGAS (Cyclic GMP-AMP synthase)-STING (Cyclic GMP-AMP/Stimulator of interferon genes) axis. cGAS represents the most extensively studied among the pattern recognition receptors (PRRs) that are involved in sensing ASFV (32–35). Mechanistically, BANF1 competes with cGAS for DNA binding both in the nucleus and the cytoplasm, thereby preventing cGAS activity on cellular self-DNA (36).

In this report, we first confirmed that BANF1 interacts with both MGF360-21R and A151R, employing Y2H, biochemical, and colocalization assays. At the functional level, an RNA interference approach was used to demonstrate the proviral effect of BANF1 in ASFV replication. Subsequently, we explored whether A151R and MGF360-21R play a role in the modulation of the IFN-α/β signaling pathway. Using different strategies, we provide unprecedented evidence for the crucial role of A151R and MGF360-21R in counteracting the induction of the IFN-α/β response and suggest that this inhibition does not involve BANF1. Finally, single A151R and MGF360-21R deletion mutants were constructed and characterized *in vitro*. Both mutant viruses were unable to efficiently control the induction of the IFN-α/β response in primary porcine macrophages compared to their parental strain. Additionally, the GeorgiaΔA151R mutant showed a growth defect in a multi-step replication kinetics assay, indicating an important role of A151R in the virus replication cycle.

## Results

### Identification of host and viral proteins interacting with ASFV MGF360-21R and A151R by AP-MS

To gain functional information about the previously uncharacterized ASFV protein MGF360-21R, we analyzed its interactions with the host and viral proteins in infected cells. To begin with, MGF360-21R of the highly virulent ASFV strain Georgia 2007/1 (37) fused to an N-terminal green fluorescent protein (GFP) tag was expressed in HEK293 cells and subsequently AP-MS was performed to identify its interactors (**Fig. 1A**). GFP alone was employed as a negative control. Interacting proteins were purified with a GFP-trap system and subjected to MS analysis in biological triplicates. We compiled a list of background proteins to filter out nonspecific binders from true protein interactors (**Supplementary Table S1A**). In this way, we selected 232 protein interactions for MGF360-21R (**Supplementary Table S1B**). The top five ranked high-confidence proteins, based on their abundance, sequence coverage, and probability score, were ASFV protein A151R, elongins B and C (ELOB and ELOC), and two pyrroline-5-carboxylate reductases (PYCR1 and PYCR2) (**Fig. 1B**). Notably, A151R emerged as both, a high-confidence interactor and the only viral protein interacting with MGF360-21R. Therefore, we also analyzed the interactome of N-terminally GFP-tagged A151R in infected HEK293 cells by AP-MS (**Supplementary Table S1C**). This allowed us to compare the interactomes of both viral proteins and to distinguish common and specific interaction patterns regulating MGF360-21R and A151R functions during viral infection. We identified 48 common interactors (**Fig. 1C**). Additionally, 184 specific interactions were identified for MGF360-21R and 33 for A151R. The gene ontology (GO) term enrichment analysis (38) and pathway enrichment analysis using the Kyoto Encyclopedia of Genes and Genomes (KEGG) database (39), or the Reactome database (40) showed that A151R interactors were significantly enriched in RNA processing and nuclear envelope reformation proteins (**Fig. 1D, Supplementary Table S1D**). The proteins interacting with MGF360-21R were enriched for terms related to mitochondrial transport (GO:1990542) and ubiquitin-dependent protein degradation (GO:0043161). Next, we focused on proteins from enriched terms and assembled a host-virus interaction network of proteins binding A151R and/or MGF360-21R (**Fig. 1E**). We used the STRING database (41) to identify interactions between host proteins that have been experimentally validated and the EBI Complex Portal (42) database to assign the proteins into specific protein complexes (**Supplementary Table S1E**).

**Figure 1.**
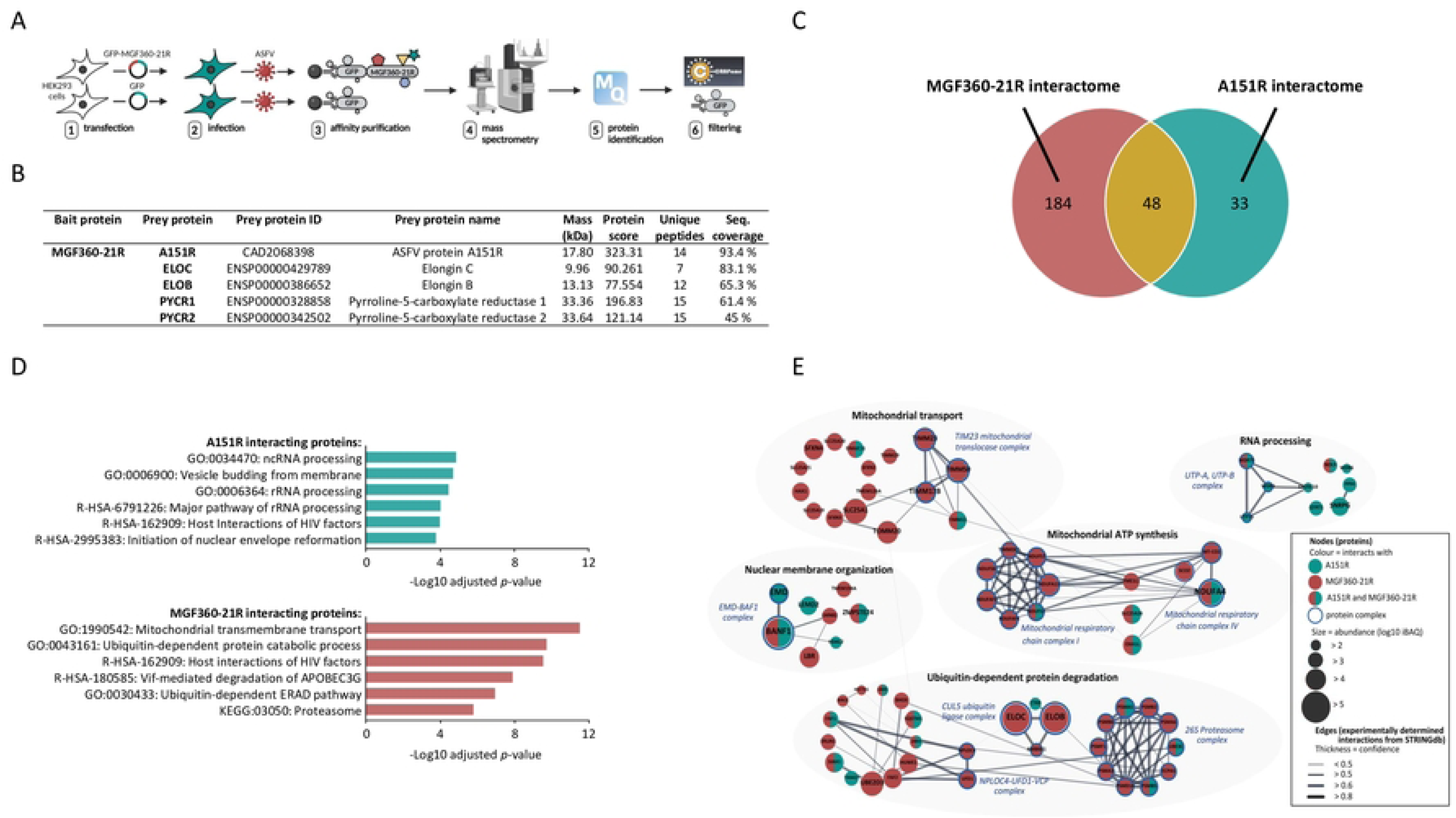
Interactome of MGF360-21R and A151R in ASFV-infected HEK293 cells. (A) AP-MS experimental workflow for identifying host and viral proteins interacting with MGF360-21R. (B) Top 5 high-confidence proteins identified as interactors for MGF360-21R in AP-MS experiments. (C) Venn diagram of proteins identified by AP-MS in ASFV-infected HEK293 cells expressing MGF360-21R-GFP or A151R-GFP. (D) Term enrichment profiles of A151R and MGF360-21R interactomes. The most significant GO, KEGG, and Reactome terms are shown with Benjamini-Hochberg corrected *p*-values. (E) Network of host proteins interacting with A151R (green nodes), MGF360-21R (red nodes), or both proteins (half green, half red nodes) clustered by enriched terms in GO, KEGG, or Reactome. Node sizes scale with protein abundances (log10 iBAQ). Protein complex constituents are indicated as nodes with blue border and the respective complex names according to the EBI Complex Portal are presented as blue descriptions. Edges indicate the protein-protein interactions based on the available experimental evidence, and edge thickness represents the confidence prediction of the interaction from the STRING database.

### Mapping cellular interactors of ASFV MGF360-21R and A151R by Y2H

Complementary to the AP-MS approach, a porcine cDNA library was screened by HT-Y2H using MGF360-21R and A151R viral proteins from Georgia 2007/1 as baits. The schematic representation of the HT-Y2H screening protocol is illustrated in **Fig. 2A**. Each screen was performed by yeast mating to obtain a minimum of 30×10^6^ diploids, a number corresponding to 10-times the complexity of our cDNA library. A total of 311 positive [His+] yeast colonies were recovered from these two screens (132 and 179 clones for MGF360-21R and A151R, respectively), and cellular prey proteins were identified by cDNA amplification, sequencing, and multi-parallel BLAST analysis. Four interactors were identified for MGF360-21R, with cytochrome c oxidase subunit 2 (COX2) and small ribosomal subunit protein uS19 (RPS15) appearing once each while TCF3 fusion partner (TFPT) was found 16 times (**Fig. 2B**). Interestingly, BANF1 emerged as a common interactor of MGF360-21R (114 times) and A151R (179 times), also being the only interactor of A151R. To confirm this result, the full-length porcine BANF1 was retested against MGF360-21R and A151R by Y2H (**Fig. 2C**). As expected, both MGF360-21R and A151R were able to interact with BANF1. Among the shared interacting proteins of MGF360-21R and A151R, BANF1 represented the highest confidence in both AP-MS and Y2H approaches. In addition, the data obtained from Y2H also indicated that BANF1 constitutes a direct binding partner of both, MGF360-21R and A151R. Therefore, we comprehensively characterized the interaction between the viral proteins MGF360-21R and A151R, along with the host protein BANF1, and explored its significance in the context of ASFV infection.

**Figure 2.**
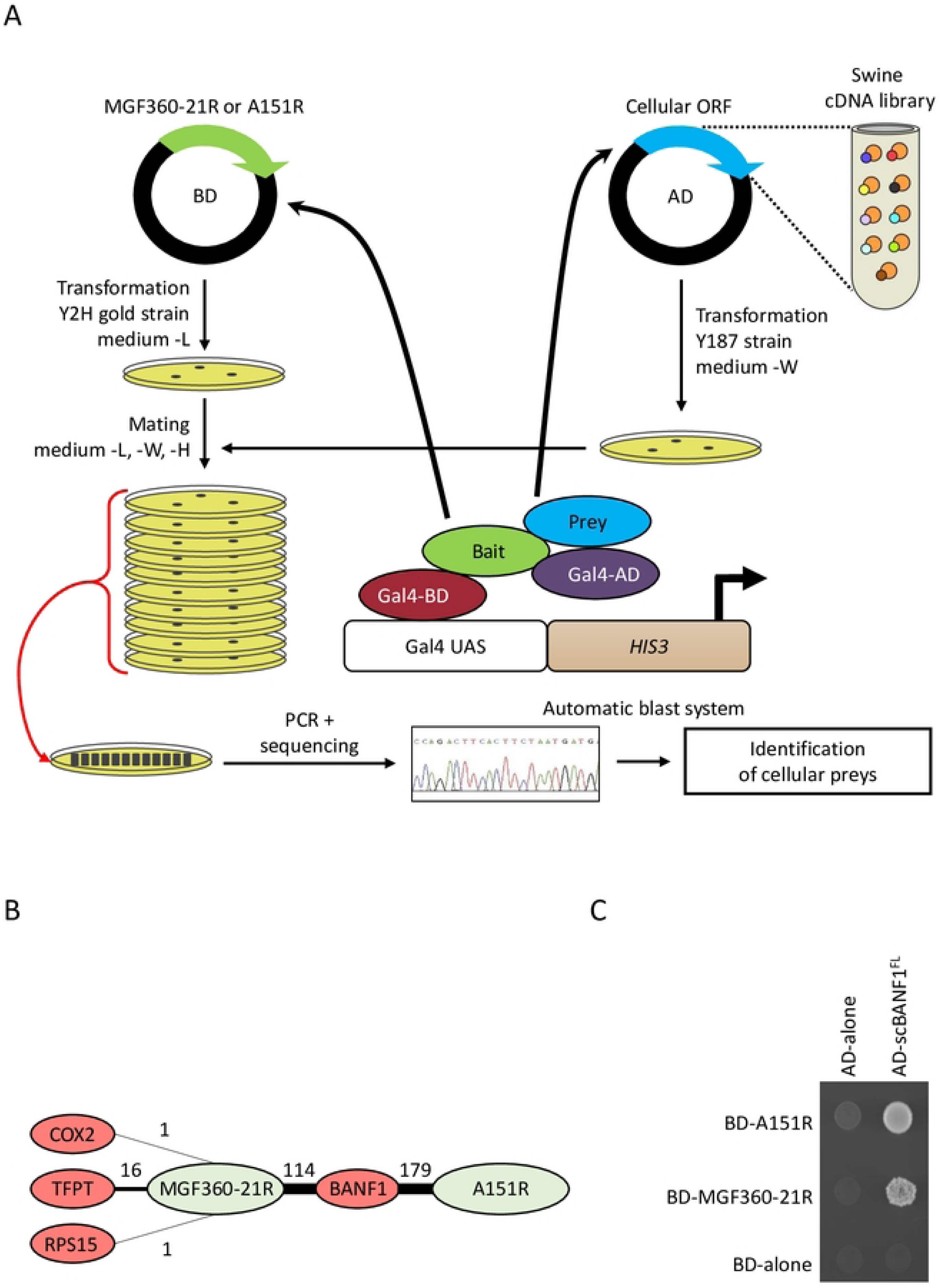
Overview of the HT-Y2H Pipeline and data. (A) A151R or MGF360-21R (viral bait) is expressed in yeast and tested for host protein interactors by mating with yeast containing libraries of swine cDNA (cellular prey). An interaction between the viral protein and the cellular protein induces transcription of a reporter gene allowing growth on selective media. Then, cDNAs from each colony are amplified by PCR, sequenced and identified by an automatic blast system. (B) For each interaction, the thickness of the edge corresponds to the number of positive yeast colonies. (C) Interactions of BANF1 with A151R and MGF360-21R were analyzed in the Y2H system. Yeast cells expressing A151R or MGF360-21R fused to the Gal4 DNA-binding domain (BD) were co-transformed with a plasmid encoding the Gal4 transactivation domain (AD) fused to BANF1. Yeast cells were plated on a growth medium supplemented with 5 mM 3-AT to test interactions.

### MGF360-21R and A151R interact with both human and swine BANF1

To validate the interactions at the biochemical level, 3×FLAG-tagged full-length swine BANF1 was co-expressed in HEK293T together with GST-tagged MGF360-21R or A151R and subsequently complexes binding to the GST-tagged ASFV proteins were affinity purified with glutathione-sepharose beads. As expected, BANF1 co-purified with both MGF360-21R and A151R (**Fig. 3A and 3B**). Moreover, after exchanging swine BANF1 with the human BANF1 in these experiments, binding of human BANF1 to A151R and MGF360-21R could also be observed (**Fig. 3C and 3D**). This validation was crucial, particularly as some of our functional studies were performed with human cells. To determine whether BANF1 directly binds to MGF360-21R or A151R and evaluate their corresponding binding affinities, 3×FLAG-tagged swine BANF1, GFP-MGF360-21R or GFP-A151R were expressed in HEK293T cells and purified using anti-FLAG or GFP affinity agarose. Subsequently, microscale thermophoresis (MST) was performed to quantify the dissociation constant (K_d_). In accordance with the previous co-immunoprecipitation data, direct binding of BANF1 to GFP-A151R was confirmed and a K_d_ value of 105±31 nM was determined for the complex (**Fig. 3E**). In contrast, a direct interaction between BANF1 and MGF360-21R could not be observed, indicating that other factors may be involved in the interaction between BANF1 and MGF360-21R in these conditions. In addition, formation of massive MGF30-21R oligomers was observed during the purification process, which might have interfered with the formation of a BANF1 – MGF360-21R complex under the conditions of MST measurement. However, structural modelling of complexes between BANF1 and MGF360-21R with varying stoichiometries using AlphaFold-Multimer resulted in a high confidence model consisting of one MGF360-21R molecule and one BANF1 dimer (43) (**Fig. 3F**) therefore we do not exclude the possibility of the formation of a MGF360-21R – BANF1 complex.

**Figure 3.**
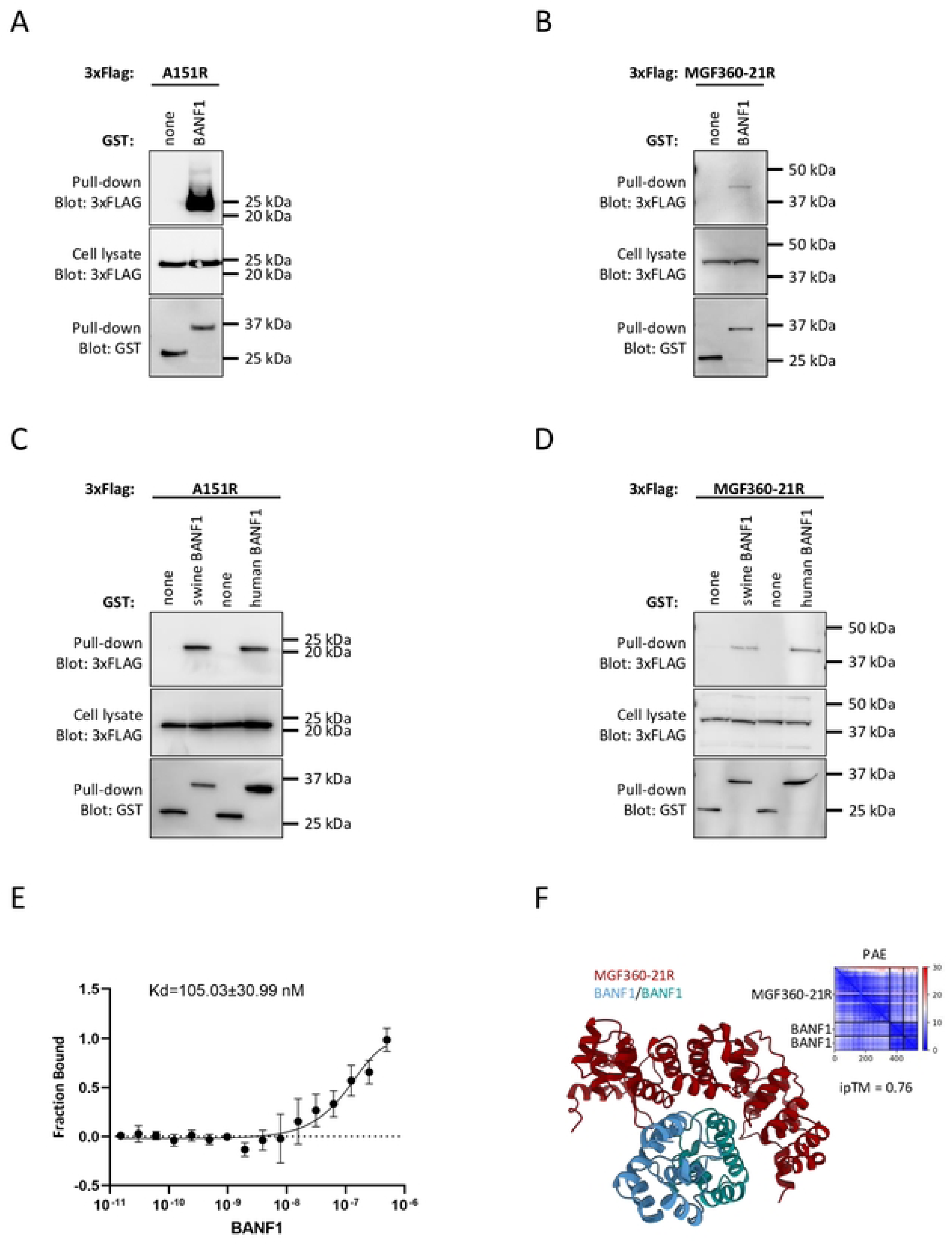
Biochemical validation of BANF1 interaction with A151R and MGF360-21R. (A-D) HEK-293T cells were transfected with expression vectors encoding GST alone or fused to either A151R (A, C) or MGF360-21R (B, D) and tested for the interaction with either human (C, D) or swine BANF1 (A-D). Total cell lysates were prepared 48 h post-transfection (cell lysate, middle panels), and co-purifications of indicated cellular proteins were assayed by pulldown using glutathione-sepharose beads (pulldown, upper panels). GST-tagged viral proteins were detected by immunoblotting using anti-GST antibody (pulldown, lower panels), while BANF1 was detected with an anti-3xFLAG antibody. Masses are shown in kilodaltons (kDa). (E) Microscale thermophoresis (MST) analysis of direct binding of A151R to BANF1. (F) Predicted structure of the MGF360-21R (red) and dimeric BANF1 (blue and green) complex with an ipTM of 0.76 as ribbon representation. Low PAE values, which indicate high confidence, between protein chains indicate potential interactions.

### A151R and MGF360-21R colocalize with BANF1 both in porcine and human cells

To assess the potential impact of the A151R-BANF1 and MGF360-21R-BANF1 interactions on their respective subcellular localizations, we first carried out fluorescence microscopy in porcine IBRS2 cells. A fusion protein of swine BANF1 with the red fluorescent protein mCherry was co-expressed with GFP-tagged A151R (GFP-A151R) or MGF360-21R (GFP-MGF360-21R) (**Fig. 4A**). When expressed alone, BANF1, MGF360-21R, and A151R were localized in the cytoplasm, with A151R also exhibiting perinuclear localization. Upon co-transfection, mCherry-tagged BANF1 colocalized with both GFP-tagged MGF360-21R and A151R in the cytoplasm and the perinuclear region. We next examined the subcellular localization of endogenous BANF1 in HEK293 cells in response to the ectopic expression of GFP-A151R, GFP-MGF360-21R, or the GFP tag alone (**Fig. 4B**). In contrast to mCherry-tagged BANF1, endogenous BANF1 was exclusively located within the nucleus when the GFP tag alone was expressed as a control. Interestingly, the expression of either A151R or MGF360-21R induced the translocation of BANF1 to the cytoplasm. Yet these two viral proteins exhibited distinct colocalization patterns with endogenous BANF1: A151R and BANF1 appeared to concentrate around the cell nucleus, while MGF360-21R and BANF1 were dispersed in the cytoplasm. Altogether, these results demonstrate that A151R and MGF360-21R colocalize with BANF1 in both porcine and human cells.

**Figure 4.**
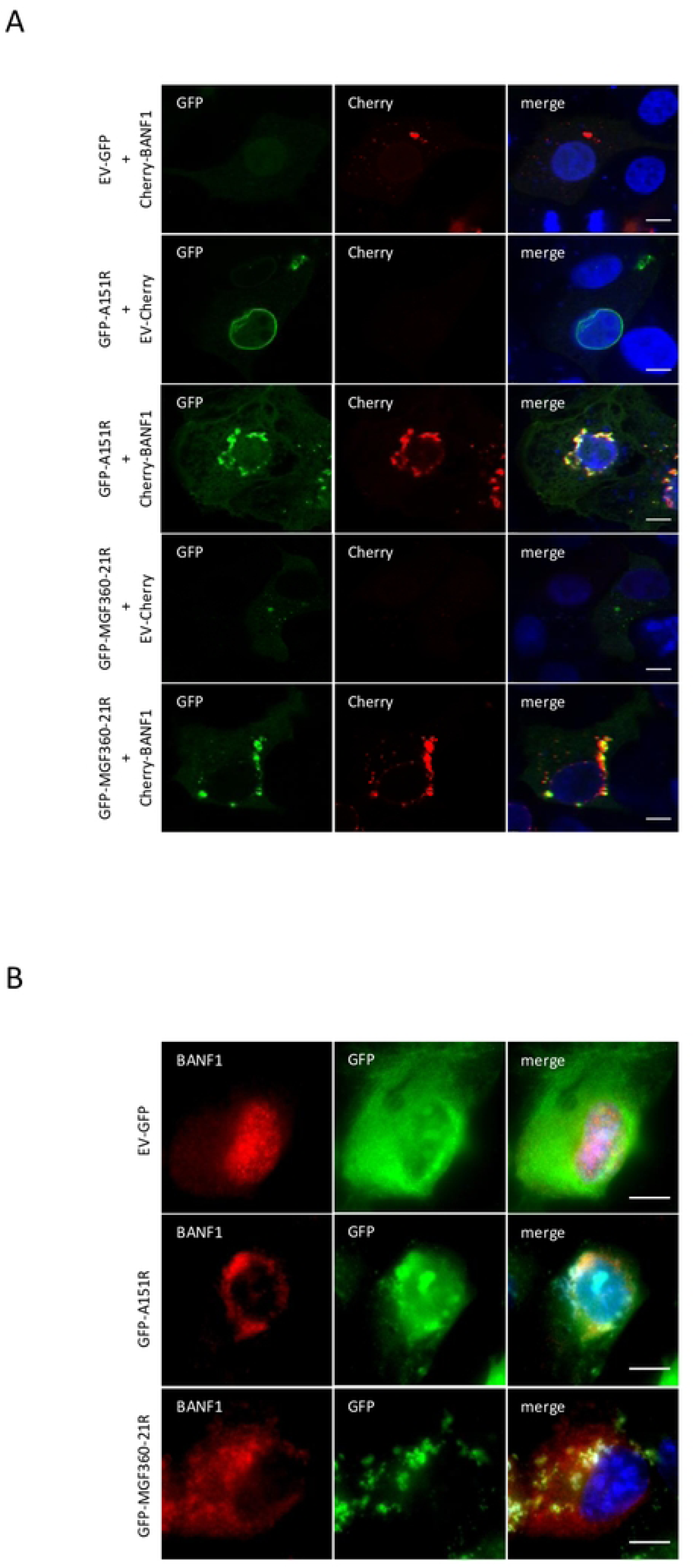
Analysis of BANF1 interactions by colocalization assays. (A) IBRS2 cells were co-transfected with pEGFP-C1 and pmCherry-C1 plasmids encoding A151R or MGF360-21R and swine BANF1, respectively. After 24 h, cells were fixed and labelled with the dye Hoechst 33258 to stain nuclei. Intracellular localization of Hoechst-stained nuclei (blue), A151R or MGF360-21R (green), and BANF1 (red) were visualized by confocal fluorescence microscopy (×40 magnification). (B) Same experiment as (A) but with HEK293 cells transiently expressing GFP-A151R and GFP-MGF360-21R to test colocalization with endogenous BANF1. Cells were visualized by confocal laser scanning microscope (×63 magnification). Scale bars represent 10 µM.

### Relocation of BANF1 during ASFV infection

As infection with vaccinia virus (VV) and Herpes Simplex Virus 1 (HSV-1) leads to marked changes in the cellular distribution of BANF1 (44,45), we have studied the relocation of BANF1 in the context of ASFV infection. Subsequently, we analyzed the subcellular localization of BANF1 in porcine WSL cells during ASFV infection (**Fig. 5A**). As expected in mock-infected cells, BANF1 was exclusively located in the nucleus. However, we failed to detect BANF1 in the cell at 8 hours (h) post infection (p.i.) or observed only minimal protein levels in the cytoplasm at 24 h p.i., suggesting ASFV infection induced BANF1 degradation or obscured its detection by antibodies. To address this question, we reanalyzed the proteomic dataset documenting changes in host protein levels during ASFV infection in porcine WSL cells generated in a prior study (46). We observed no significant changes of BANF1 protein abundance throughout the course of the infection (**Fig. 5B**), indicating that BANF1 is not undergoing ASFV-induced degradation.

**Figure 5.**
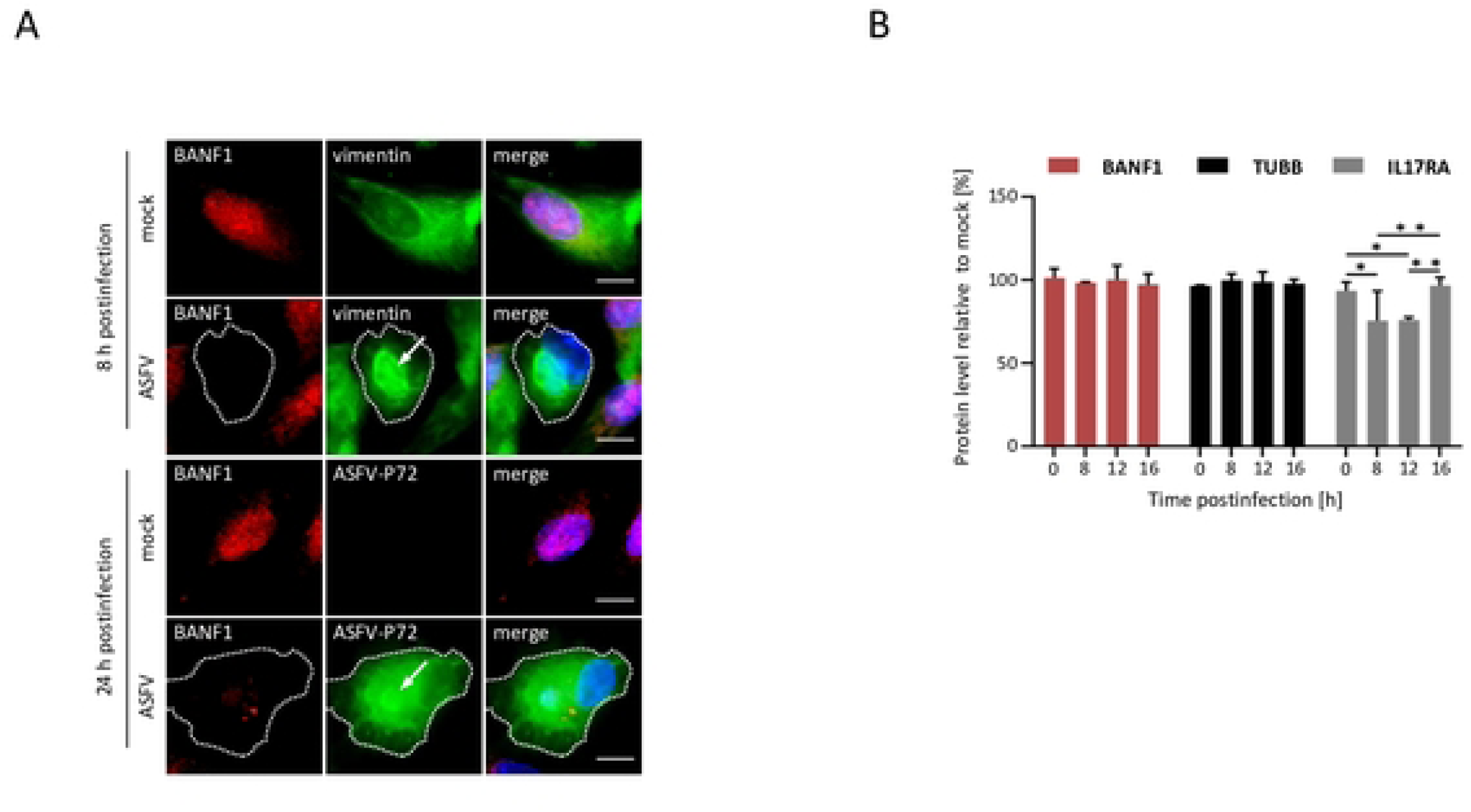
Alterations in subcellular localization and abundance of BANF1 during ASFV infection. (A) Indirect immunofluorescence shows BANF1 distribution in mock and ASFV-infected WSL cells monitored at 8 and 24 h p.i.. Virus replication sites (virus factories), marked with arrows, were labeled with antibodies directed against vimentin (at 8h p.i.) and against ASFV-P72 protein (at 24 h p.i.). Dotted lines indicate the outlines of infected cells, bars represent 10 μm. (B) Comparison of BANF1 protein levels in mock and ASFV-infected WSL cells based on log10 LFQ (label-free quantitation) values. Levels of tubulin beta chain (TUBB) and interleukin-17 receptor A (IL17RA) are presented as controls. Means of results from three independent replicates and standard deviations are plotted. *, *p* < 0.05; **, *p* < 0.01.

### BANF1 silencing reduces ASFV replication

Given that BANF1 has been demonstrated to function as a retroviral cofactor (47,48) while also displaying antiviral properties against VV (49) and HSV-1 (45), we decided to examine its influence on ASFV replication in the porcine WSL cell line. For this purpose, BANF1 expression was knocked down by short interfering RNA (siRNA) pools (si-BANF1). In parallel, we used a nonspecific siRNA (si-NegC) pool as a control. Titration of si-BANF1 ranging from 1 nM to 10 nM revealed effective silencing of BANF1 in WSL cells at concentrations of 3 nM and 6 nM, as confirmed through Western blotting analyses (**Fig. 6A**) and immunofluorescence microscopy (**Fig. 6B**). Importantly, the suppression of BANF1 expression in WSL cells did not decrease cell viability when compared to si-NegC (**Fig. 6C**). Next, WSL cells were transfected with either 3 nM or 6 nM si-BANF1 for 24 h, inoculated with ASFV at a multiplicity of infection (MOI) of 1 and the infection was allowed to proceed for 0, 24, or 48 h. The progeny virus production was reduced by approximately 2-log_10_ after 24 and 48 h p.i. when using a 6 nM concentration of si-BANF1, in contrast to control cells treated with si-NegC and non-transfected (NT) cells (**Fig. 6D**). We therefore conclude that BANF1 is required for the unimpaired replication of ASFV in WSL cells. To detect any secondary effect of BANF1 downregulation on the inhibition of ASFV production induced, for instance, by impacting the host cell proteome, we performed a comparative proteomic analysis of WSL cells after treatment with si-BANF1 and si-NegC control, respectively. Of the 203 differentially expressed proteins, 92 were exclusively identified in si-BANF1 knockdown cells, and 37 exclusively in si-NegC cells (**Fig. 6E**, **Supplementary Table S2**). Additionally, 67 proteins were significantly upregulated, and 7 significantly downregulated in si-BANF1 treated cells. The suppression of BANF1 led to increased expression of proteins involved in the virus-mediated innate immune response through cGAS-STING, interferon, and RIG-I-like receptor pathways (**Fig. 6F**). BANF1 knockdown also affected the expression of proteins participating in DNA and histone methylation, DNA replication, and nuclear transport. Moreover, the Ser/Thr kinase VRK1 (vaccinia-related kinase 1), responsible for the phosphorylation of BANF1 (49), was only identified in siBANF1 knockdown cells. We also detected changes in the levels of emerin complex proteins (MYOE1, MYH9, ACTB) and zinc metalloproteinase Ste24 homolog (ZMPSTE24), directly implicated in the functioning of BANF1 binding partners, emerin, and lamin A/C (50,51).

**Figure 6.**
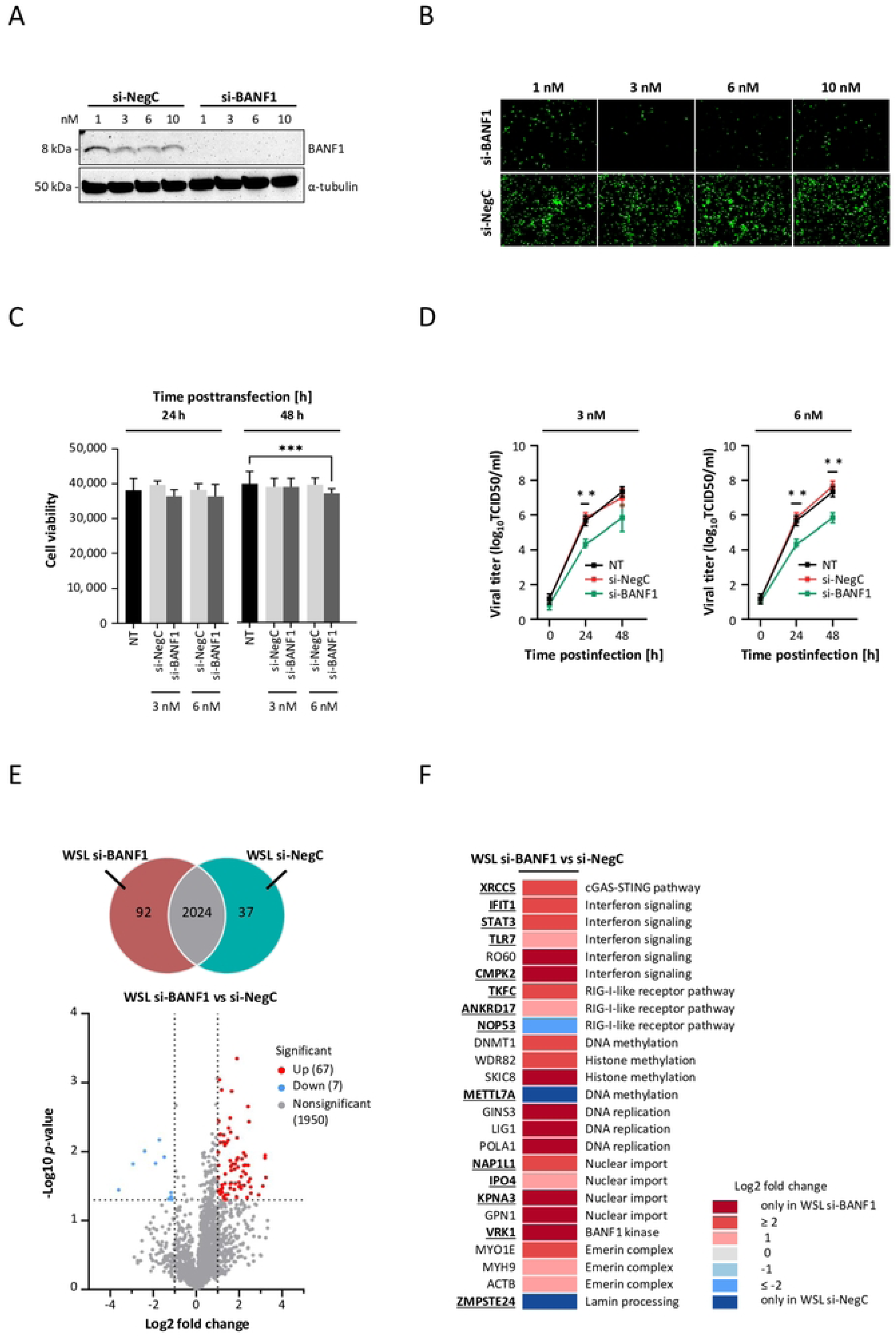
The effect of BANF1 downregulation on ASFV replication and the cellular proteome. (A) Expression of BANF1 in WSL cells transfected with increasing concentrations of si-BANF1 or si-NegC detected by Western blot analyses. The α-tubulin-specific signal was used as a loading control. (B) Immunofluorescence images show endogenous BANF1 in WSL cells transfected with increasing concentrations of si-BANF1 and si-NegC. (C) Cell viability of WSL cells without treatment (NT) or after treatment with si-NegC, or si-BANF1 at the given concentrations was quantitated with a resazurin-based assay (PrestoBlue). Corrected *p*-values below 0.001 are indicated by ***. (D) Growth kinetics of ASFV on WSL cells without treatment (NT) or after incubation with si-NegC and si-BANF1 after infection at a MOI of 1 (n = 3 wells/cell line/time point). The culture medium was collected at the indicated times, and the yields of the cell-free virus were expressed as TCID50 per milliliter and were plotted as means of results from three independent replicates and standard deviations, ** indicates *p* < 0.01. (E) Venn diagram of proteins identified by MS in WSL si-NegC and si-BANF1 knockdown cells (upper panel). Volcano plot showing differentially abundant proteins in WSL si-NegC and si-BANF1 knockdown cells (bottom panel). The −log10 *p*-value (Benjamini– Hochberg corrected) is plotted against the log2 (fold change: si-BANF1/si-NegC). The dotted vertical lines denote ±1.0-fold change on the log2 scale while the dotted horizontal line denotes the significance threshold of *p* = 0.05. (F) Heatmap showing a selection of proteins characterized by significant changes in abundance in WSL cells after si-NegC and si-BANF1 knockdown. Proteins known to be involved in virus replication (see references in **Supplementary Table S2**) have been highlighted in bold and are underlined.

### A151R and MGF360-21R inhibit the induction of the IFN-α/β signaling pathway

Based on our findings presented in **Fig. 6F**, and considering the established role of BANF1 in the regulation of the cGAS/STING-dependent IFN response, we hypothesized that A151R and MGF360-21R expression might inhibit this pathway. To answer this question, an IFN-β reporter assay was used. HEK293T cells were co-transfected with a plasmid encoding the luciferase gene under the control of the IFN-β promoter. This could be activated by the plasmid-driven expression of cGAS and STING. The impact of A151R or MGF360-21R on IFN-β expression was measured by co-expression of the viral proteins together with the IFN-β promoter luciferase reporter system and the cGAS/STING inducers. A151R and MGF360-21R efficiently reduced the stimulating effect of cGAS/STING (**Fig 7A**). To further assess whether the observed inhibitions were specific to the cGAS/STING axis, we performed the same experiments using cells stimulated with poly(dA:dT), which mimics viral double-stranded DNA, or overexpressed with a constitutively active mutant of RIG-I (NΔRIG-I, N-terminal CARDs of RIG-I). As shown in **Fig. 7B** and **7C**, similar inhibitions of IFN-β promoter activity were observed for both A151R and MGF360-21R. However, neither of the two ASFV proteins antagonized an IFN-stimulated response element (ISRE)-luciferase gene reporter (**Fig. 7D**) in IFN-β-stimulated cells, indicating that they do not inhibit Jak/STAT signaling and that the antagonism is specific to the IFN induction pathway. Although HEK293T cells are highly efficient for transfection, we aimed to complement our analysis by carrying out equivalent experiments using porcine cells. To do so, we measured the expression level of *IFN-*β mRNAs in porcine PK15 cells treated by either the cytosolic DNA sensor ligands: interferon stimulated DNA (ISD) or poly(dA:dT). As shown in **Fig. 7E** and **7F**, we confirmed that A151R and MGF360-21R could also modulate the induction of the IFN-α/β signaling pathway in a porcine *in vitro* model. It should be noted that A151R showed a very strong inhibition of this cellular pathway following ISD stimulation (**Fig. 7E**). To investigate if the inhibition of IFN-α/β activation by A151R and MGF360-21R was dependent on their interaction with BANF1, we used a gene silencing approach. HEK293T cells were transfected with BANF1-specific or control nonspecific siRNA before transfection with 3×FLAG-tagged A151R or MGF360-21R and stimulation by cGAS/STING or poly(dA:dT) (**Fig. 7G** and **7H**, respectively). In both cases, the reduction of BANF1 expression did not affect the capacity of A151R and MGF360-21R to control the IFN-β induction. Under these conditions, we conclude that binding to BANF1 may not represent a molecular mechanism underlying the inhibition of the IFN-α/β pathway by A151R and MGF360-21R.

**Figure 7.**
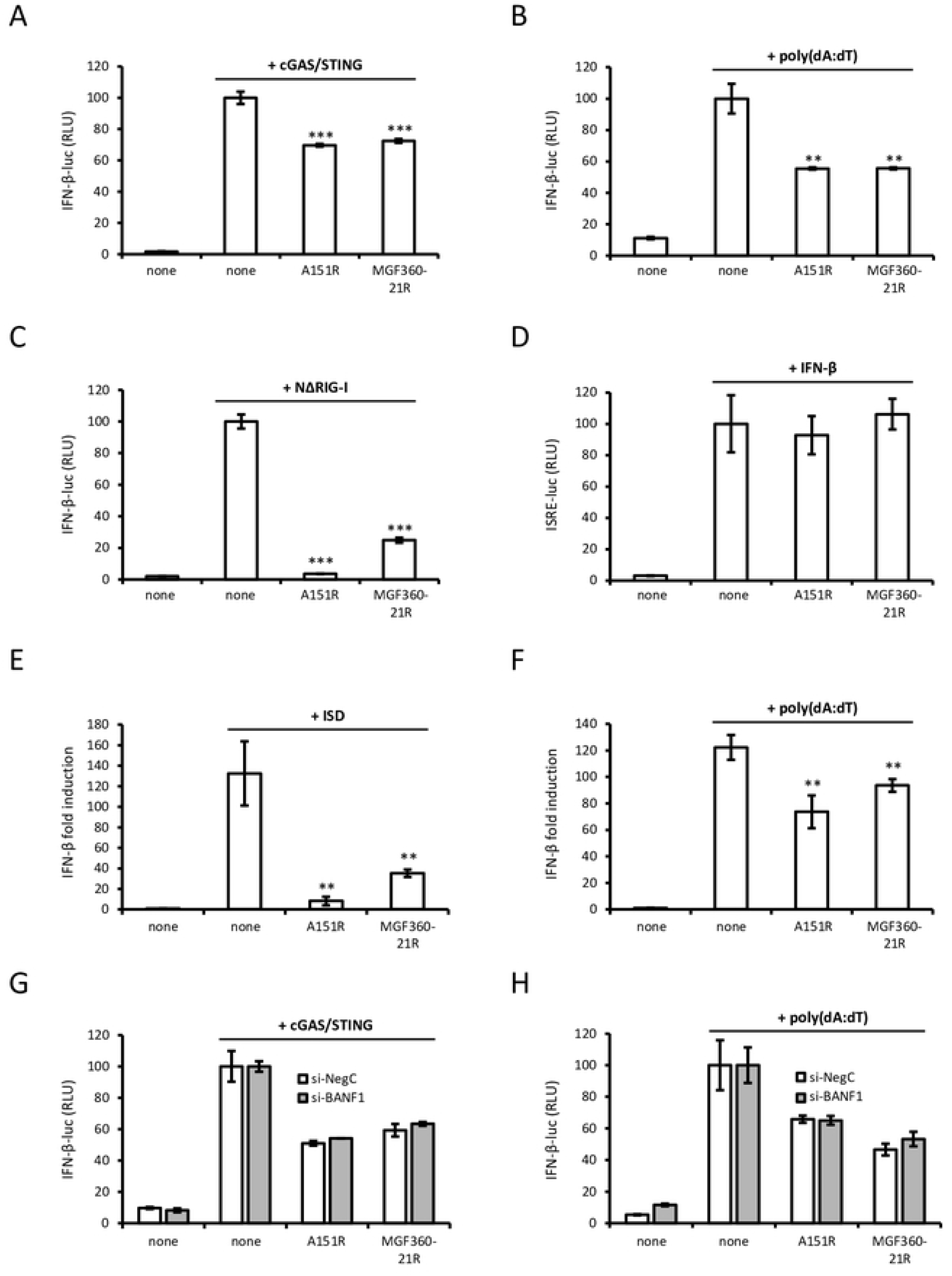
Activation of the IFN-β and ISRE promoters in cells expressing A151R and MGF360-21R proteins. (A) HEK293T cells were co-transfected with IFN-β-pGL3 and pRL-CMV reference plasmids, pUNO1 plasmids expressing cGAS and STING and pCI-neo-3×FLAG expression vectors encoding 3×FLAG alone or fused to A151R or MGF360-21R. After 48 h, the relative luciferase activity was determined. (B, C) Same experiment as (A) but with HEK293T cells transfected with poly(dA:dT) (B) or with a plasmid encoding NΔRIG-I (C). (D) Same experiment as (A) except that HEK293T cells were stimulated after 24 h with 1000 IU/ml of IFN-β and expression of the luciferase reporter construct controlled by ISRE repeats (pISRE-Luc) was quantified 24 h later. (E, F) PK15 cells were transfected with either the cytosolic DNA sensor ligand ISD (E) or poly(dA:dT) (F) together with pCI-neo-3×FLAG expression vectors encoding 3×FLAG alone or fused to A151R or MGF360-21R. After 36h, the expression level of *IFN-β* was measured by RT-qPCR and normalized to that of *GAPDH*. Data are presented as a fold increase relative to the non-stimulated condition. (G, H) Same conditions as (A, B) but with HEK293T cells knockdown BANF1 (si-BANF1) and control cells (si-NegC). All experiments were achieved in triplicate. Data represent means ± SD and are representative of three independent experiments. **, p < 0.005 and ***, p < 0.0005.

### BANF1 interactome in ASFV-infected cells

Beyond a role in innate immunity, BANF1 facilitates various essential cellular processes by engaging with DNA, histones, and many other nuclear and cytoplasmic proteins (reviewed in reference (31)). To investigate the BANF1 interactome in the context of ASFV infection and DNA binding, we expressed the BANF1-GFP fusion protein in WSL cells under the four following experimental conditions: (i) absence of infection and benzonase treatment (mock-infected, no treatment), (ii) absence of infection and presence of benzonase treatment (mock-infected, benzonase treatment), (iii) presence of ASFV infection and absence of benzonase treatment (ASFV-infected, no treatment), and (iv) presence of ASFV infection and benzonase treatment (ASFV-infected, benzonase treatment). Similar to the previous AP-MS experiment, we compiled a list of background proteins to eliminate nonspecific binders (**Supplementary Table S3A**). We compared the protein identifications from the four datasets and identified several BANF1-protein interactions that were solely detected in either mock-infected or ASFV-infected cells (**Fig. 8A**). Notably, the number of DNA-dependent interactions unique to BANF1 was comparatively lower in infected cells than in non-infected cells. Term enrichment analysis of the proteins exclusively detected in mock-infected and infected cells highlighted their involvement in distinct cellular processes: regulation of chromatin remodeling (GO:0006338) and DNA replication (GO:0006260) in non-infected cells and nuclear transport (GO:0051168) and mRNA splicing (GO:0000398) in ASFV-infected cells (**Fig. 8B** and **Supplementary Table S3B**). Furthermore, we constructed a heatmap representing the BANF1 interactomes across various conditions (**Fig. 8C** and **Supplementary Table S3C**). A significant proportion of the high-confidence binding proteins play a role in diverse functions such as chromatin remodeling, transcription regulation, nuclear membrane organization, nuclear transport, and innate immune responses. Among these proteins, host factors that facilitate or hinder viral replication were present. In contrast to several ASFV proteins which co-purified with BANF1 exclusively in the presence of DNA, the interaction between BANF1 and A151R was detected even in the absence of DNA, confirming that DNA is not required and it is indeed a direct protein-protein interaction (PPI). Next, we concentrated on identifying the presence of established BANF1 interactors and protein complexes within our dataset (**Fig. 8D** and **Supplementary Table S3D**). Our data demonstrated that many interactions previously described between human BANF1 and other human proteins are also present in porcine cells. Additionally, in ASFV-infected cells, BANF1 interacted with actin-like protein 6A (ACTL6A) and histone H1.3. Conversely, the transcription factor POU2F1 and the serine/threonine-protein kinase (VRK2), responsible for phosphorylating BANF1, were exclusively identified in the absence of ASFV infection. Of the proteins not previously reported as BANF1 interactors, we noticed that those associated with various chromatin remodeling complexes and NFκB signaling pathway (RELA and NFκB1) were primarily identified in mock-infected samples. Moreover, RELA and NFκB1 were only detected in presence of DNA, suggesting that BANF1 could be recruited to the NFκB-responsive promoters and this complex is likely disrupted by ASFV. The binding of MDA5 and TLR7 indicates that BANF1 would also modulate IFN-I induction downstream of PRRs other than cGAS/STING. Interestingly, two components of the TREX transcription-export complex, the spliceosome RNA helicase (DDX39B) and THO complex subunit 7 (THOC7), were exclusively identified in pulldowns from infected samples. Notably, the TREX complex was reported to be essential for exporting Kaposi’s sarcoma-associated herpesvirus mRNA and virus replication (52).

**Figure 8.**
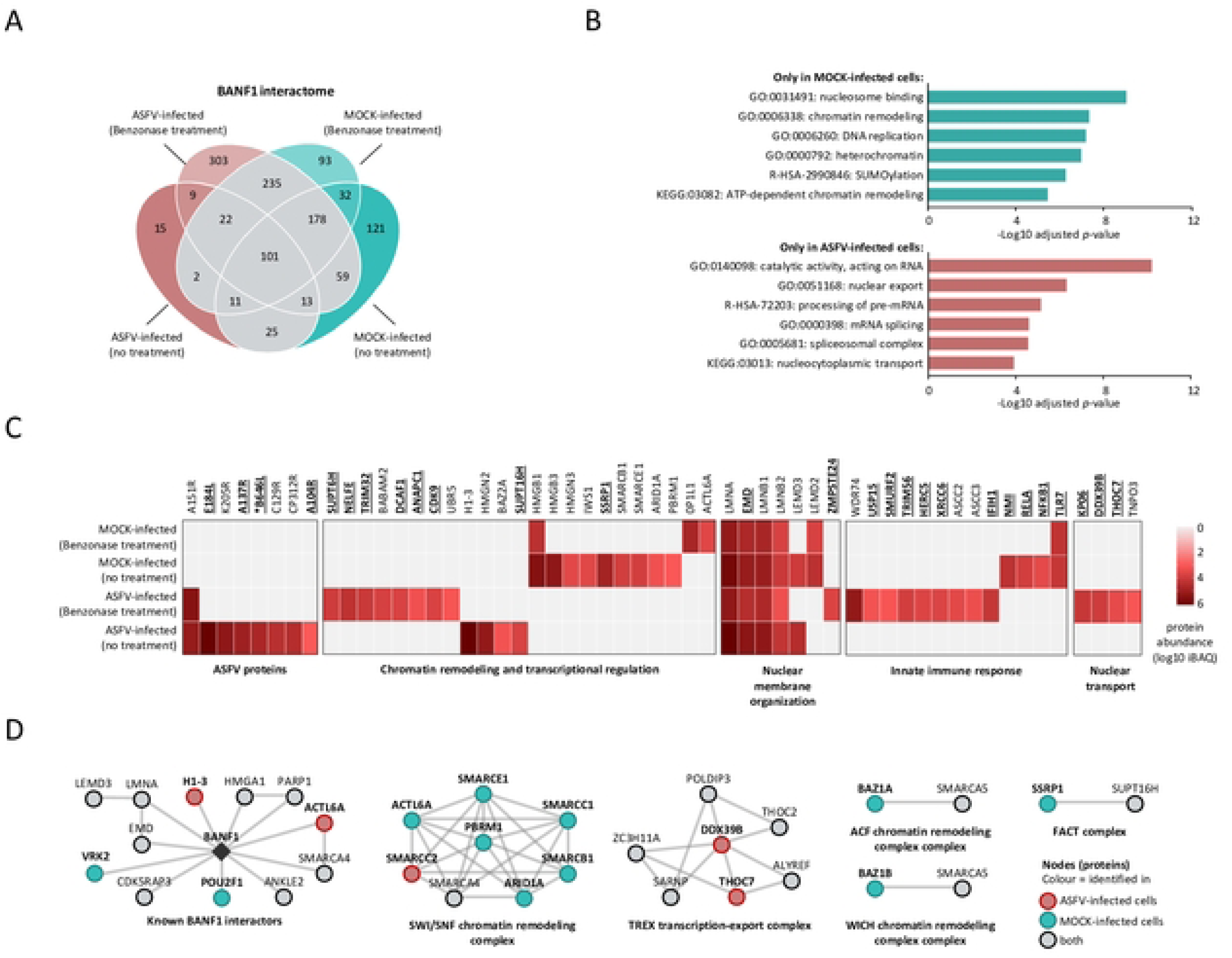
Comparison of the interactomes of BANF1 prepared from non-infected and ASFV-infected cells. (A) Venn diagram of proteins identified by AP-MS in WSL cells expressing BANF1-GFP, categorized by combinations of the infection status (mock or ASFV) and benzonase treatment (no or 25 U/mL benzonase). (B) Enrichment profiles of proteins interacting with BANF1 in mock-infected and ASFV-infected samples. The most significant GO, KEGG, and Reactome terms are shown with Benjamini-Hochberg FDR-corrected p-values. (C) Heatmap of high-confidence BANF1 interactors compared under different experimental conditions. Functional characterizations for grouped proteins are shown below the heatmap. Proteins known to be involved in virus replication (see references in **Supplementary Table S3C**) are highlighted in bold and underlined. ASFV proteins that are essential for virus replication are marked with an asterisk (*) (D) The high-confidence proteins that bind to BANF1 were grouped into protein complexes according to the EBI Complex Portal and visualized with STRING to illustrate the experimentally validated protein-protein interaction among them. Node colors indicate the identification in ASFV-infected (red), mock-infected (green), or both (grey nodes) samples.

### Generation and characterization of ASFV GeorgiaΔA151R and GeorgiaΔMGF360-21R deletion mutants

In order to further understand the role of A151R and MGF360-21R in the context of the viral infection of primary macrophages, two viruses with these single genes deleted (GeorgiaΔA151R and GeorgiaΔMGF360-21R) were purified as described in materials and methods. Full genome sequencing confirmed that the deletions occurred at the expected genome positions: 49,653 to 50,107 for GeorgiaΔA151R and 187,834 to 188,782 for GeorgiaΔMGF360-21R (**Fig. 9A and 9B**). The later deletion leaves 267 bp of the MGF360-21R gene remaining (**Fig. 9B**), which could lead to the translation of a truncated protein. Indeed, upstream of the second ATG in the remaining sequence there is a transcription start site that would produce a 61 amino acid protein (53). However, this protein is very unlikely to retain the function of the full length MGF360-21R. With the exception of a single T to A point mutation in the E199L gene that leads to a E125V mutation in pE199L in the GeorgiaΔA151R virus, no other mutations were observed as compared to the wild type virus. The ability of the recombinant viruses to replicate *in vitro,* was assessed over a multi-step growth curve (**Fig. 9C)**. Porcine bone marrow cells (PBMCs) were infected with the deletion mutants or the wild type virus at a MOI of 0.01. Total virus from both cells and supernatants were collected at different times p.i. and titrated. All the viruses reached a plateau around 72 h p.i. with maximum titers of 10^7.25^ hemeadsorbing dose 50% HAD_50_/ml for WT and GeorgiaΔMGF360-21R and 10^6.38^ HAD_50_/ml for GeorgiaΔA151R. Hence, the GeorgiaΔA151R mutant showed a growth defect, and this was most noticeable at 48 h p.i. (p = 0.0009). Given the impact of both proteins on the induction of type I IFN in transfected cells (**Fig. 7**), we evaluated the secretion of IFNα (**Fig. 9D**) and CXCL-10 (**Fig. 9E**), an interferon stimulated gene (ISG), by PBMCs infected with WT or recombinant viruses. At 8 h post infection, supernatants from GeorgiaΔA151R infected cells contained significantly higher amounts of IFNα (p = 0.0175). At 16 h post infection, all supernatants contained substantial amounts of IFNα and again these were higher in GeorgiaΔA151R infected cells (p < 0.0001). A similar trend was observed for CXCL-10. At both 8 and 16 h post infection, CXCL-10 levels were significantly higher following infection with GeorgiaΔA151R (p < 0.0001). Notably, cells infected with GeorgiaΔMGF360-21R also secreted higher amounts of CXCL-10 than those infected with the wild type virus at 16h p.i. (p = 0.0028). This indicates that MGF360-21R might have a smaller, but still significant, impact on the host IFN response.

**Figure 9.**
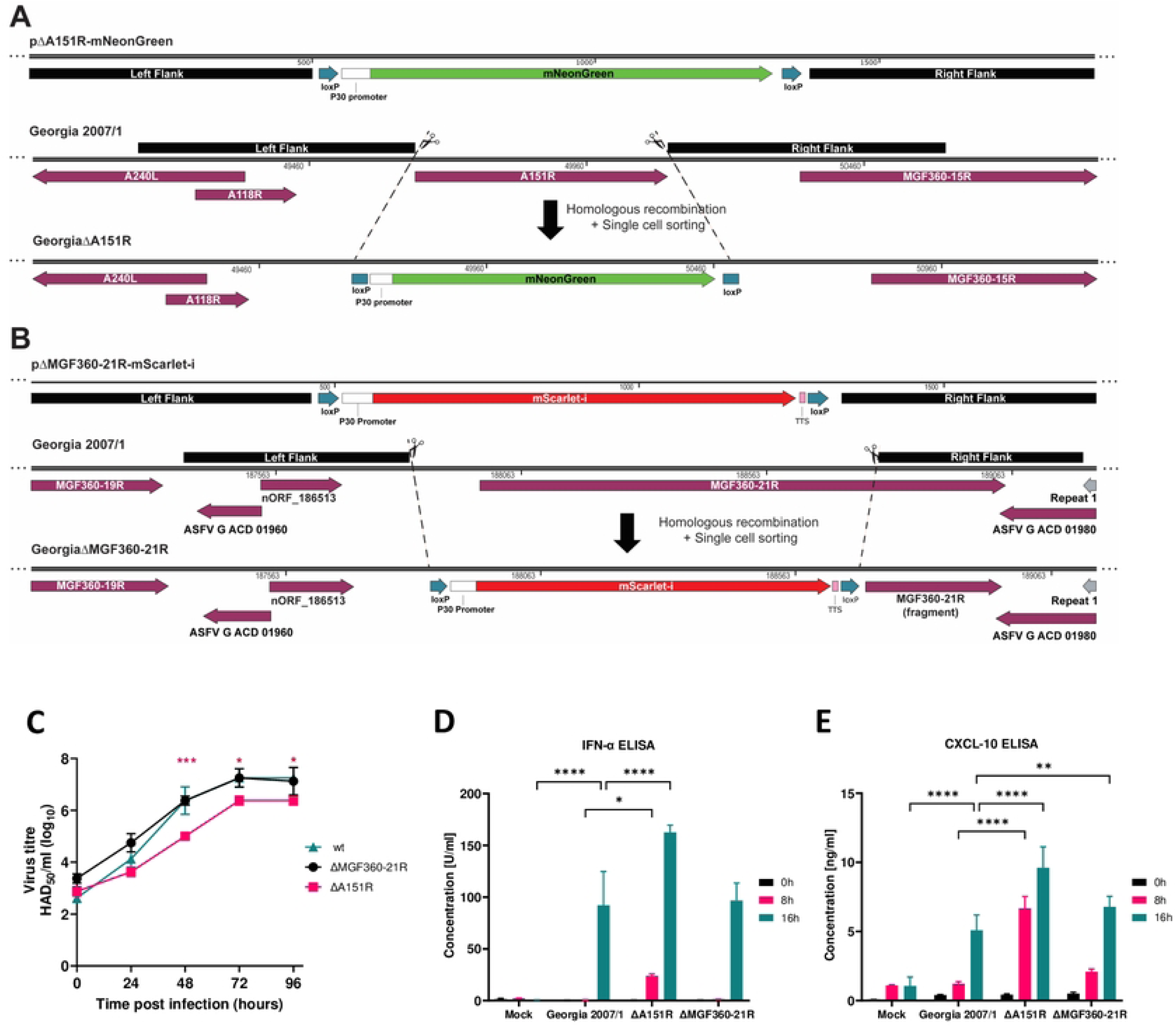
Purification, replication and IFN responses of ASFV GeorgiaΔA151R and GeorgiaΔMGF360-21R deletion mutants. (A, B) Diagram showing plasmid constructs, sites of homologous recombination and final sequence of GeorgiaΔA151R and GeorgiaΔMGF360-21R deletion mutants respectively. (C) Multistep growth curves of the WT and gene-deleted viruses over a course of four days, where day 0 represents the inoculum. Means and standard deviations from two independent titrations are shown. Significant differences are represented by asterisks, where * is p < 0.05 and *** is p < 0.001. (D, E) Levels of IFN-α (D) and CXCL10 (E) from supernatants of purified PBMs infected with WT or recombinant viruses or mock infected. Means and standard deviations from duplicate measurements are shown. Significant differences, as compared with supernatants from wild type Georgia/2007 virus infected cells, are represented by asterisks, where * is p< 0.05, ** is p < 0.01 and **** is p< 0.0001.

## Discussion

Identifying virus–virus and virus–host PPIs provides important insights into molecular mechanisms of virus replication and pathogenesis. Over the past two decades, many novel PPIs have been reported for ASFV proteins (reviewed in reference (4)). However, about half of the currently annotated ASFV proteins still await characterization while the full panel of ASFV-coded proteins may not be completely determined yet (1,53). Discovering new molecular virus-host interactions provides information about the functionality of so far uncharacterized viral proteins and cellular pathways involved in viral infection and pathogenesis and provides a basis for the rational development of antiviral drug targets and vaccines.

The starting point for this study was to investigate the potential functions of MGF360-21R, an uncharacterized protein within the ASFV MGF360. Previous research has shown that MGF360 and MGF505 members act as multifunctional immune-evasion proteins. They inhibit type I IFN responses by interacting with key proteins in the cGAS-STING (14,15,54–58) and JAK-STAT (59–61) signaling pathways, often leading to their degradation. These results, however, are primarily based on targeted PPI studies focused on host proteins linked to immunity. This approach introduces a bias and leaves the specific host proteins, involved in regulation of other important pathways, unidentified. Here, we applied AP-MS and Y2H, as ‘open-view’ approaches, for the less biased identification of MGF360-21R interactors from the full complement of host and virus proteins present in an infected cell. Interestingly, MGF360-21R was co-purified alongside another viral protein, A151R, which has recently been recognized to inhibit IFN-β production (62,63). Interactions between viral proteins can diversify targets inside the host cell and collectively influence its internal environment. Next, we focused on identifying potential interactions involving viral proteins MGF360-21R and A151R with host proteins.

We identified BANF1 as the high-confidence mutual interactor of ASFV MGF360-21R and A151R using AP-MS (**Fig. 1E**) and Y2H screening (**Fig. 2B**). The observed interactions were subsequently validated through co-immunoprecipitation (**Fig. 3**) and colocalization studies (**Fig. 4**). Furthermore, MST experiments substantiated that the interaction between A151R and BANF1 is indeed direct (**Fig. 3**), which is an important finding considering that BANF1 is a known DNA-binding protein. We also demonstrated that the expression of either A151R or MGF360-21R leads to a relocalization of endogenous BANF1 from the nucleus to the cytoplasm. However, we were unable to detect the subcellular localization of BANF1 in the context of early ASFV infection despite no apparent degradation had been observed based on MS data. Thus, detection of BANF1 may have failed because critical epitopes detected by the anti-BANF1 monoclonal antibody might have been masked by post-translational modifications induced on BANF1 in response to the infection, thereby preventing recognition by the antibody. It is known that VV and HSV-1 infections can modulate the phosphorylation state of BANF1 (44,45,64,65). Phosphorylation at the threonine and serine residues at positions 2, 3, and 4 by VKR kinases regulate BANF1 function (66,67), particularly its DNA binding affinity (26,45,68), and localization (49). In our studies we showed that the expression level of VRK1 is significantly increased in the absence of BANF1 (**Fig. 6F**), further investigations are warranted to determine whether ASFV could potentially influence the phosphorylation of BANF1 and its role in ASFV replication.

BANF1 is targeted by some viruses, which hijack its function for their replication. In our report, we demonstrated that BANF1 silencing led to greatly reduced viral titers suggesting a beneficial role of BANF1 in AFSV replication. The proviral function of BANF1 has already been described in the context of infection by various retroviruses such as HIV (Human Immunodeficiency Virus) or MoMLV (Moloney Murine Leukemia Virus), with BANF1 being part of their pre-integration complex (47,69). In Gammaherpesviruses, including KSHV (Kaposi’s Sarcoma-Associated Herpesvirus) and EBV (Epstein-Barr Virus), BANF1 facilitates lytic reactivation by inhibiting cGAS-mediated DNA sensing (70). Conversely, BANF1 can also exert an antiviral function in the replication of VV (64) and HSV-1 (45). This is achieved through its ability to bind their viral genome, thereby inhibiting genome replication.

BANF1 was originally discovered and named for its role in protecting retroviral DNA against suicidal autointegration (48). The DNA binding properties allow BANF1 to recognize and bind to dsDNA of both foreign (47,71) and endogenous origin (24). Within the cytoplasm, BANF1 was shown to compete for DNA binding with cGAS (36), consequently restricting the cGAS-STING pathway and suppressing innate immune responses (72). These facts prompted us to investigate whether the interaction of MGF360-21R and A151R with BANF1 could regulate type I IFN induction during ASFV infection. While the expression of MGF360-21R and A151R have no effect on ISRE-luciferase gene expression in IFN-β-stimulated cells, they significantly inhibit the induction of IFN-α/β signaling downstream of cGAS/STING or RIG-I. Our findings correlate with a recent study showing the inhibitory effect of A151R on IFN-I induction (62,63). Moreover, a A151R deletion mutant of the Georgia 2010 ASFV strain exhibits reduced virulence in domestic pigs and induces a protective response against experimental infection with its parental virulent strain (19). In our experiments, we have also shown that macrophages infected with a modified Georgia 2007/1 strain lacking A151R exhibit higher levels of IFN-α at 8h and 16h p.i. compared to those infected with the parental virulent Georgia 2007/1 strain (**Fig. 9**). Our findings not only confirm the crucial role of A151R in inhibiting the IFN-I pathway but also support previous observations showing similar significant difference in IFN-α/β production between avirulent/low virulent and high virulent ASFV strains (7,35,73–75). In contrast, the anti-IFN-I activity of MGF360-21R was revealed for the first time in this study, adding to other ASFV proteins belonging to the MGF genes family (including MGF505-11R, MGF360-13L, MGF505-7R, MGF360-11L, MGF505-3R, MGF110-9L, MGF360-14L, MGF360-15R and MGF505-2R) which have already been shown to inhibit IFN-I induction (32,34,76–80). These genes have evolved through homologous recombination (81), which seems to be an effective mechanism employed by ASFV to generate genetic diversity and develop strategies to evade host immunity (5). Poxviruses seem to have evolved a similar mechanism, enabling them to adapt and to bypass host defenses despite their low mutation rate (82).

Although both MGF360-21R and A151R can impact IFN-I induction, this inhibition may occur independently of their interaction with BANF1. Conversely, suppression of BANF1 expression in porcine cells increases the expression of proteins linked to the innate immune response activation through the cGAS-STING and RIG-I pathway (**Fig. 6F**) and significantly reduces ASFV replication (**Fig. 6D**). Therefore, we cannot exclude a role of BANF1 in suppressing innate immune defenses. However, if BANF1 does not directly contribute to the inhibition of IFN-I by MGF360-21R and A151R, the question remains regarding the mechanism underlying this inhibition. Furthermore, we suggest that ASFV A151R could downregulate the IFN-I signaling by acting as a transcriptional regulator, potentially suppressing the transcription of IFNs themselves and downstream IFN-regulated genes. During infection, protein A151R is expressed in both the early and late stages (83) and has been observed to accumulate within the nucleus and the nuclear envelope (**Fig. 4**). The crystal structure of A151R (18) revealed a Zn-binding site typically linked to interactions with DNA, RNA, and with various proteins (84). We hypothesize that A151R might modulate gene transcription through direct or indirect interactions with nucleic acids. Like EBV’s Zta protein (85), A151R could recognize and bind directly to specific DNA sequences within the host genome. On the other hand, many viral transcriptional regulators only bind DNA when in complex with host transcription factors. This characteristic could also apply to A151R.

As more functional studies appear, it becomes clear that ASFV proteins have the potential to display moonlighting characteristics, meaning their functions can vary due to alterations in cellular localization, infection stages, cell types, or changes in the concentration of a cellular or viral ligand. We demonstrated the inhibitory role of ASFV proteins MGF360-21R and A151R in IFN-I signaling. Nonetheless, both proteins might perform additional functions attributed to their interaction with BANF1. Given the high expression of A151R and MGF360-21R within 2 to 6 h p.i., it is highly likely that both proteins have substantial roles in the early phase of infection. Previous studies indicate that within the initial four hours of infection, ASFV triggers the formation of nuclear blebs (86) and disrupts the lamina network, releasing nuclear material into the cytoplasm (87). The nuclear lamina is a filamentous meshwork forming an interface between the inner nuclear membrane and the peripheral chromatin. BANF1 is an integral component of the lamina network, which is essential for nuclear envelope assembly (88). Therefore, further investigations are needed to assess whether the interaction with BANF1 (i) contributes to the disruption of the lamina network caused by ASFV, and (ii) facilitates the transport of viral DNA and proteins between the nucleus and cytoplasm, enhancing viral replication.

## Materials and methods

### Cell culture

The HEK293 and the wild boar lung-derived (WSL) cells (89) were supplied by the Friedrich-Loeffler-Institut Biobank (catalog numbers CCLV-RIE 0197 and 0379, respectively). Cells were maintained in Iscove’s modified Dulbecco’s medium (IMDM) mixed with Ham’s F-12 nutrient mix (1:1 [vol/vol]) supplemented with 10% fetal bovine serum (FBS). HEK293T and PK15 cells were maintained in DMEM + Glutamax medium (Fisher) containing 10% FBS (Eurobio), 1 mM sodium pyruvate (Fisher), 1% penicillin-streptomycin (Fisher) and 1% non-essential amino acids (Fisher). IBRS2 cells were maintained in MEM medium (Fisher) supplemented with 1.5% lactalbumin hydrolysate (Sigma), 7% FBS (Eurobio), 1% penicillin-streptomycin (Fisher) and 2.5% Hepes (Fisher). PBMCs were obtained from the leg bones of four-to five-week-old, outbred pigs. Following density gradient centrifugation, using Histopaque-1083, the mononuclear cell fraction was recovered, washed, and maintained in Roswell Park Memorial Institute (RPMI) supplemented with 10% FBS, 1% penicillin-streptomycin, and 100 ng/mL porcine colony-stimulating factor CSF1 (Roslin Tech). All cell types were maintained at 37°C with 5% CO2.

### Plasmid DNA constructs

A151R and MGF360-21R of the Georgia 2007/1 strain were cloned by gene synthesis (Twist Bioscience) into pTWIST-ENTR plasmid. Swine-BANF1 and human-BANF1 were amplified from PMA and A549 cDNA libraries respectively, using specific primers flanked with the Gateway cloning sites 5′-GGGGACAACTTTGTACAAAAAAGTTGGC and 5′-GGGGACAACTTTGTACAAGAAAGTTGG. PCR products were cloned by *in vitro* recombination into pDONR207 (Gateway System; Invitrogen). ORFs were then transferred from pTWIST-ENTR or pDONR207 into different Gateway-compatible destination vectors, according to the manufacturer’s recommendations (LR cloning reaction; Invitrogen) and their sequences were verified (Eurofins). GST, Gal4-BD, GFP, mCherry and 3×FLAG tag fusions were achieved using pDEST27 (Invitrogen), pDEST32 (Invitrogen), pDEST-EmGFP-vivid colors, pmCherry-C1 or pCI-neo-3×FLAG vector, respectively.

Transfer plasmids for generation of ASFV recombinants were synthesized (Genscript, UK) to express the fluorescent markers mNeonGreen or mScarlet-I under the control of the ASFV P30 promoter, for the deletion of A151R or MGF360-21R respectively. The markers’ sequences were flanked by around 500 bp of the left and right flanking regions of the genes to be deleted (**Fig. 9A and Fig. 9B**).

### Antibodies

The primary antibodies used for immunofluorescence were mouse anti-vimentin (MA1-06908, Thermo Fisher), rabbit anti-BANF1 (ab129184, Abcam). Additionally, a rabbit antiserum specific for ASFV B646L (P72) (90) was used at dilution of 1:20,000 for immunoblotting. The secondary antibodies were Alexa Fluor 647-conjugated goat anti-rabbit IgG (H+L) (A21245, Invitrogen) and goat anti-mouse IgG (H+L) (A32728, Invitrogen). The primary antibodies used for immunoblotting included mouse anti-tubulin (B-5-1-2; Sigma-Aldrich).

### Transfection

HEK293 cells were transiently transfected with a GFP-MGF360-21R, a GFP-A151R, a GFP-BANF1, or a GFP vector using a K2 multiplier and K2 transfection reagent (Biontex) following the manufacturer’s instructions. At 24 h post-transfection, cells were infected with ASFV. For each bait, three independent biological replicates were prepared for affinity purification.

### ASFV infection in WSL cells

All experiments with ASFV were performed in a biocontainment facility, fulfilling the safety requirements for ASF laboratories and animal facilities (Commission Decision 2003/422/EC, chapter VIII). ASFV (Armenia/07 isolate) was adapted by serial passaging to more efficient replication in WSL cells. Passage 20 stocks were generated as described previously (91) and used in infection experiments. Cell monolayers were inoculated with ASFV stock dilutions at an MOI of 1 PFU/cell, and supernatants collected from mock-infected cells were used as controls. After inoculation, cells were centrifuged for 1 h at 600 × g and 37°C. Next, cells were washed three times with phosphate-buffered saline (PBS), replenished with medium containing 5% FBS, and incubated at 37°C with 5% CO_2_. Supernatants were harvested at appropriate times, and progeny virus titers were determined as 50% tissue culture infective doses (TCID_50_) per milliliter (92) on WSL cells.

### Affinity purification and mass spectrometry

#### Affinity purification

A total of 5 × 10^6^ cells were seeded for each AP experiment. After an overnight incubation, cells were transiently transfected for 24 h before infection with ASFV. At 24 h post-infection and 48 h post-transfection, cells were washed, lysed, and affinity purified on 50 μL GFP-trap agarose beads (Chromotek) as described previously (46).

#### On-bead digestion

Bead-bound proteins were suspended in 300 μL freshly made UA buffer (8 M urea, 100 mM Tris-HCl, pH 8.0), loaded onto 10-kDa filter units (Sartorius), and centrifuged at 12,000 × g at 20°C for 30 min. Filter-aided sample preparation (FASP) trypsin digestion was performed as described previously (93). Proteins were trypsinized on beads in 100 μL of digestion buffer (1 M urea, 50 mM Tris-HCl, pH 7.5, and 5 μg/mL trypsin (V5111, Promega)). Digestion was performed overnight at 37°C with shaking. The next day, the peptide containing supernatant was collected by ultrafiltration and acidified with formic acid (1% final concentration). Peptides were desalted using C18 100-μL tips (Thermo Scientific) according to the manufacturer’s instructions, dried by vacuum centrifugation, and reconstituted in 20 μL of 0.1% formic acid before mass spectrometry.

#### MS data acquisition and analysis

Samples were analyzed on a timsTOF Pro mass spectrometer coupled to a nanoElute nanoflow liquid chromatography system (Bruker Daltonics). Peptides were separated on a reversed-phase analytical column (10 cm × 75 μm i.d., Bruker 1866154) by application of a binary gradient made from 0.1% formic acid (FA) in water (solvent A) and 0.1% FA in acetonitrile (solvent B). During chromatography solvent B was raised over 60 min (2% to 4% from 0 to 1 min, 4% to 20% from 1 to 46 min and 20% to 32% from 46 to 60 min) at a constant flow rate of 250 nL/min. The column temperature was maintained at 40°C. MS analysis of eluting peptides was performed in ddaPASEF mode (1.1-s cycle time) as recommended by the manufacturer. Proteomic data were searched against an ENSEMBL *Homo sapiens* proteome (v.GRCh38.p13) or a *Sus scrofa* proteome (v.11.1.2021-11-10) and an NCBI ASFV Georgia (v.FR682468.2) proteome database using the default settings for MaxQuant (v.2.0.3.0) (94). The false discovery rates (FDR) on the peptide and protein levels were set to 1%, the minimum peptide length was seven amino acids, and the match-between-runs option was used with a 0.7-min match window and 20-min alignment time. The output files were analyzed using Perseus software (v.2.0.10.0) (95). A protein was considered as identified if at least 2 unique peptides were found in 2 out of 3 of the replicates. Proteins specifically binding to MGF360-21R, A151R, or BANF1 baits were filtered out by removing the GFP background. The background list consists of proteins identified in our GFP negative controls and proteins identified as common protein contaminants for AP-MS experiments in HEK293 cells and deposited in the Contaminant Repository for Affinity Purification (CRAPome) (96). Potential interactors were considered specific if they were identified only in GFP-bait pulldowns or if the log2 fold change (between GFP control and GFP-bait) was greater than 2 and the *p*-value of a two-sided t-test was <0.01.

#### Data availability

All MS raw data and MaxQuant output tables were deposited in the ProteomeXchange Consortium (http://proteomecentral.proteomexchange.org) via the PRIDE partner repository (97) and will be publicly available upon final publication (identifier PXD047794; accession for reviewers: Username; Password: HdEHCqNb).

### Term enrichment analysis and interaction network

Porcine genes corresponding to the identified proteins were assigned to their corresponding human orthologues using the R package gprofiler2 (v.0.2.1) (98). The interactors of each bait were tested for enrichment of GO biological processes, KEGG annotations, and Reactome terms. The overrepresentation analysis was performed using the enricher function of the clusterProfiler (v.4.2.2) (99) package in R with default parameters. Selected proteins from significantly enriched functional groups were manually curated, and the network diagram was plotted using Cytoscape (v.3.7.2) (100) with the Cytoscape StringApp plugin (101).

### Yeast two-hybrid (Y2H) screening procedure

Our Y2H protocol closely followed the methodology previously described (102). ORFs encoding A151R and MGF360-21R were fused with a Gal4 DNA-binding domain (Gal4-BD), within the pDEST32 vector (Invitrogen). These constructs were transformed into Y2H Gold yeast strain (Clontech) and selected on a selective medium lacking leucine (-L). Y2H screens were performed using a swine cDNA library, which has been generated with mRNAs extracted from porcine alveolar macrophages (PAM) and then cloned into the Gal4-AD pDEST22 vector (Creative Biogene). For each screen, at least 30 million yeast diploids were produced and grown on selective medium lacking leucine, tryptophan and histidine (-L, -W, -H) and supplemented with 5 mM of 3-aminotriazole (3-AT). After 6 days, yeast colonies were picked and purified over 3 weeks by culture on selective medium -L, -W, -H + 5 mM of 3-AT to eliminate false-positives (103). Yeast colonies were lysed using zymolyase (Euromedex) and AD-cDNAs were amplified by PCR. Those PCR products were sequenced (Eurofins), and cellular preys were identified by a multiparallel BLAST analysis.

### GST pulldown experiments

HEK293T cells were plated in a 6-well plate at a density of 2 × 10^6^ cells/well and transfected 24 h later (JetPRIME, Polyplus) with 500ng of pDEST21 encoding GST-BANF1 (swine or human) and 3x-flag-tagged A151R or MGF360-21R expressed in pCI-neo-3xFLAG. After 36 h, cells were harvested in PBS and incubated on ice cold-lysis buffer (20 mM MOPS-KOH pH 7.4, 120 mM KCl, 0.5% Igepal, 2 mM β-Mercaptoethanol), supplemented with Complete Protease Inhibitor Cocktail (Roche), for 20 min. The cell lysates were then clarified at 14,000×g for 30 min. Protein extracts were incubated for 2 h at 4 °C on a spinning wheel with 30 μL of glutathione-sepharose beads (Amersham Biosciences). The beads were washed three times for 5 min with lysis buffer on a spinning wheel, and samples were boiled in denaturing loading buffer (Invitrogen).

### Microscale thermophoresis analysis

The binding of A151R and MGF360-21R to BANF1 was measured by microscale thermophoresis (MST). After purification, the concentration of GFP-A151R was measured with the fluorescence-based method by normalizing the fluorescence intensity of GFP-A151R to the FITC standard curves as described before (104). 20 nM GFP or GFP-A151R proteins in MST buffer (50 mM Tris-HCl, pH 7.5, 150 mM NaCl, 10 mM DTT) were incubated with different concentrations of ligands. Immediately, samples were loaded into standard glass capillaries (NanoTemper) and thermophoresis analysis was performed on a NanoTemper Monolith NT.115 instrument (40% LED, 80% MST power) at 22°C. A laser on-time of 30 s and a laser off-time of 5 s were used. The experiment was performed in triplicates, and the MST curves were fitted using NT analysis software to obtain the Kd values.

### Immunoblotting

Samples were heated for 5 min at 95°C and resolved by SDS-polyacrylamide gel electrophoresis (SDS-PAGE) on 4 to 20% Mini-Protean TGX gels (Bio-Rad) (105) and transferred to the nitrocellulose membrane by semidry transfer (Trans-Blot Turbo; Bio-Rad Laboratories) (106). All membranes were blocked in 5% milk powder in Tris-buffered saline with 0.25% Tween-20 (TBST) and probed for a minimum of 1h at room temperature with the indicated primary antibodies using appropriate dilutions. GST and 3×FLAG-tagged proteins were detected with a rabbit polyclonal anti-GST antibody (1:2500, Sigma-Aldrich) and a mouse monoclonal HRP-conjugated anti-FLAG antibody (M2 1:10000; Sigma-Aldrich), respectively. Next, membranes were incubated with peroxidase-conjugated secondary antibodies diluted in TBST. Protein bands were detected using the Clarity Western enhanced chemiluminescence (ECL) substrate (Bio-Rad), imaged on a C-DiGit blot scanner (LI-COR), and analyzed with Image Studio software (v.5.2).

### Colocalization assays

IBRS2 cells were plated in 24-well plates (Ibidi) with 1× 10^5^ cells/well. 24 h later, cells were transfected with either 250 ng of N-EmGFP-DEST Vector expressing either A151R or MGF360-21R and 250 ng of pmCherry-C1 encoding swine_BANF1. 24 h post-transfection, cells were fixed using a 4% paraformaldehyde (PFA) solution (Electron Microscopy Sciences) for 30 min and treated with PBS-glycine (0.1 M) for 5 min. DNA was stained with a Hoechst 33342 dye (1/10,000) (Life technologies) for 30 min. Finally, cells were visualized using a Leica DMI 8 confocal microscope (×40 magnification).

### Immunofluorescence microscopy

Coverslips with cells were fixed with 3.7% formaldehyde in PBS at room temperature for 60 min. Following fixation, cells were permeabilized with 0.01% TritonX-100 in PBS for 15 min and then blocked with PBS containing 10% FBS for 1 h. Coverslips were incubated with the primary antibody for 1 h at room temperature. Following washing with PBS, cells were incubated with a secondary antibody for 1 h at room temperature. Coverslips were then washed in PBS, and DNA was stained for 15 min with 1 μg/mL Hoechst 33258 (Sigma-Aldrich). Coverslips were mounted on glass slides using ProLong Glass Antifade Mountant (Invitrogen). Single-slice fluorescence images were acquired on a Leica DMI6000 TCS SP5 confocal laser scanning microscope (63× objective) and were processed with ImageJ software (v.1.52a) (107).

### Gene silencing by siRNA

Pooled siRNA against porcine or human BANF1 was custom synthesized and purchased from siTOOLs Biotech together with a nonspecific siRNA negative control. Transfections of siRNA were performed with Lipofectamine RNAiMAX (Thermo Fisher Scientific) following the manufacturer’s instructions. The proteome changes after BANF1 knockdown were analyzed by LC-MS/MS analysis. Lysates of WSL knockdown BANF1 (si-BANF1) and control cells (si-NegC) (100 μg) were prepared using the Thermo EasyPep Mini MS sample preparation kit (Thermo Scientific) according to the manufacturer’s instructions. Dried peptides were reconstituted in 0.1% formic acid to a final concentration of 100 ng/μL. Peptides corresponding to 200 ng protein were measured by LC-MS/MS and analyzed as described in “MS data acquisition and analysis.“

### Luciferase reporter gene assays

HEK293T cells were plated in 24-well plates with 5 × 10^5^ cells/well. After 24 h, cells were transfected (jetPRIME, Polyplus) with either 3×-FLAG-tagged A151R or MGF360-21R along with 300 ng of IFN-b-pGL3 or pISRE-Luc plasmid (0.3 mg/well, Stratagene) that contain the firefly luciferase reporter gene downstream of an IFN-β-specific promoter sequence or the ISRE enhancer element, respectively. Cells were also co-transfected with the normalization pRL-CMV plasmid (0.03 mg/well, Promega) as well with plasmids encoding indicated proteins (RIG-I or cGAS/STING) to stimulate the IFN-β-specific promoter. When specified, cells were transfected 24 h later with 0.1 mg/well of poly(dA:dT) (Invivogen) or treated with 1 × 10^3^ IU/ml of recombinant IFN-β (PBL Assay Science). After 48 h post-tranfection, cells were lysed (Passive lysis buffer, Promega), and both firefly and Renilla luciferase activities were detected using the Bright-Glo and Renilla-Glo luciferase assay system, respectively (Promega). Luminescence was measured using the GloMax plate reader (Promega). All graphs depict the mean ratios between luciferase and Renilla of triplicate samples and error bars of the standard deviation were calculated using Prism 7, version 7.0.

### Assessment of IFNβ levels by RT-qPCR

PK15 cells were plated in 24 wells plates with 5×10^5^ cells/well. 24 h later, cells were transfected (jetPRIME, Polyplus) with either 300 ng of 3×-FLAG-tagged A151R or MGF360-21R and, when indicated, with 1 μg of interferon stimulatory DNA (ISD, Invivogen) or 100 ng poly(dA:dT) (Invivogen). After 36 h, total RNAs were extracted (RNAeasy kit, Qiagen) RT-qPCR assays were performed using the QuantiNova SYBR Green RT-PCR kit (Qiagen) to measure the expression of the swine *IFN-β* gene. The data were then analyzed using the 2ΔΔCt method, where the amount of target, normalized to the endogenous reference *GAPDH* gene and relative to an experimental control. The results are expressed as relative fold change (Fc) in comparison with the non-stimulated condition.

### Computational modelling of protein complexes

Protein complex predictions were generated using the ColabFold version 1.5.5 (108) implementation of AlphaFold-Multimer (109). Various complex stoichiometries between MGF360-21: BANF1 (1:1 to 2:2) and A151R:BANF1 (1:1 to 5:2) were predicted and the model confidences assessed. A total of 5 models were predicted for each candidate complex with 3 recycles. Model relaxation and energy minimization was performed using the integrated Amber module. The confidence of resulting protein complex predictions was assessed based on the predicted aligned error (PAE) scores using an in-house Python script available at https://github.com/QuantitativeVirology/AlphaFold_Analysis, file score_colabfold_pairwise.py. For each pairwise combination in a multi-polypeptide prediction, the relevant portion of the PAE plot was extracted. It was evaluated by two criteria: 30 (the maximum value) minus the minimum of the PAE portion (to measure the most confident value) and percent standard deviation (to measure how broad the distribution of confidence values are, *i.e.* if there are both high confidence and low regions).

### Purification of Georgia/2007 recombinant viruses

Recombinant gene deleted ASFV were produced by homologous recombination followed by single cell sorting. Briefly, WSL cells were infected with Georgia/2007 WT virus and transfected with the transfer plasmids described above. Single cells expressing either mNeonGreen (ΔA151R) or mScarlet-I (ΔMGF360-21R) were isolated via fluorescence-activated cell sorting (FACS) into purified PBMs as previously described (110). After three rounds of single cell sorting and two rounds of limiting dilutions, viral DNA was extracted using MagVet universal isolation kit (Thermo Fisher Scientific), and the KingFisher flex extraction system (Thermo Fisher Scientific). Deletion of target genes and the absence of parental virus was confirmed by PCR using appropriate internal primers. Full genome sequencing of the recombinants was done was done as previously described using an Illumina MiSeq instrument (111). Raw reads are available at the Sequence Read Archive in BioProject <u>PRJNA1183819</u>.

### Multistep growth curve

PBMs were seeded at 4×10^5^ cells per well and infected at MOI of 0.01. Both cells and supernatants were collected at the indicated time points and freeze-thawed twice. Debris were pelleted by centrifugation and titrations were performed using PBMs from two different pigs. Virus titrations were carried out in quadruplicate by hemadsorption assay (HAD_50_/ml) and titers were calculated using the Spearman and Kärber algorithm. Two-way analysis of variance (ANOVA) followed by Dunnett’s multiple-comparison test was used to evaluate the differences between titers at different times post infection.

### Quantification of IFN-α and CXCL10 levels in supernatants

Purified PBMs from an outbred pig were seeded at 1×10^6^ cells/mL and infected with recombinant or wild-type viruses at a MOI of 0.5. After 1 h incubation the inoculum was removed and replaced with fresh medium. The supernatants were collected at 8 and 16 h post infection and centrifuged at 300 × g for 5 min to remove cells and debris. The levels of IFN-α in these supernatants were then evaluated using an in-house ELISA. Briefly, Maxisorp plates (Nunc) were coated with anti-pig IFN-α antibody (clone K9) at 0.5 µg/mL in 0.05 M coating buffer overnight at room temperature. Plates were washed with PBS-T (0.05% Tween 20 in PBS) and blocked with 1% bovine serum albumin (BSA) in PBS. Standards (recombinant porcine IFN-α, PBL Assay Science) and samples were then added in duplicate and incubated at room temperature for 2 h. Following washing, biotinylated anti-pig IFN-α antibody (clone F17) diluted 1:5000 in blocking buffer was added to the wells and incubated for 2 h at room temperature. The plates were then washed, incubated with streptavidin-horseradish peroxidase (R&D Systems, DY998), and finally developed with 3,3′,5,5′-tetramethylbenzidine (TMB) substrate (R&D Systems, DY999). After stopping the reaction with 2 N H_2_SO_4_, the absorbances were read at 450 nm. The concentration of CXCL10 in the supernatants was quantified using the swine CXCL10 Do-It-Yourself ELISA (Kingfisher Biotech). Briefly, Maxisorp plates were coated overnight with capture antibody (anti-swine CXCL10 polyclonal antibody, PB0119S-100) at 2.5 μg/mL in PBS. The plates were then washed with PBS-T and blocked with 4% BSA for 1 hour at room temperature. Samples and standards were then added, and the plates were incubated at room temperature for 1 hour. After four washes with PBS-T, detection antibody (biotinylated anti-swine CXCL10 polyclonal antibody, PBB1138S-050), diluted in blocking buffer at 0.05 μg/mL, was added to plates and incubated for another hour at room temperature. The plates were then washed, incubated with streptavidin-horseradish peroxidase and developed as described above. Two-way analysis ANOVA followed by Dunnett’s multiple-comparison test was used to evaluate the differences between IFNα and CXCL-10 concentrations in the supernatants at different times post infection.

## Acknowledgments

We thank the Biobank of the Friedrich-Loeffler-Institut for providing the WSL cell line. We also thank Barbara Bettin for her technical assistance.

